# Mutual synchronization pattern as a complementary indicator of the short-term blood pressure – heart rate feedback regulation activity

**DOI:** 10.1101/397208

**Authors:** Nikita S Pyko, Svetlana A Pyko, Oleg A Markelov, Oleg V Mamontov, Mikhail I Bogachev

## Abstract

We suggest a complementary indicator of the blood pressure – heart rate feedback regulation based on their synchronization pattern assessed by Hilbert transform. We determine the synchronization coefficient *Sync* as the fraction of time fragments where the standard deviation of the differences between instantaneous phases of blood pressure and pulse intervals are below a certain threshold. While *BRS* characterizes the *intensity* of the pulse intervals response to the blood pressure changes during observed feedback responses, the *Sync* likely indicates how often such responses are *activated* in the first place. Data from 95 tilt test records indicate that in both healthy subjects and patients with moderate autonomic dysfunction *BRS* and *Sync* are typically reciprocal suggesting that *low intensity* of the feedback responses characterized by low *BRS* is rather compensated by their *more frequent activation* indicated by higher *Sync*. In contrast, in diabetes patients with autonomic neuropathy *BRS* and *Sync* are positively correlated likely indicating the breakdown of this compensation in some of the diabetic patients. Therefore we suggest that *Sync* could be used as an additional indicator of the blood pressure – heart rate feedback regulation activity that is complementary to the widely used baroreflex sensitivity (*BRS*).

## 1. Introduction

Blood pressure levels are being simultaneously affected by multiple internal and external factors that require continuous activity of several regulatory mechanisms to keep its levels within a certain homeostatic range. Among multiple mechanisms that are involved in the regulation of blood pressure, arterial baroreceptor reflex or simply baroreflex appears one of the key mechanisms that govern short-term feedback responses to various physical stresses such as exercise, adaptation to the changes in the body position, reactions to drugs, changes in the subject’s mental conditions and so on. The increase of blood pressure is sensed by baroreceptors located in blood vessels that in turn invoke a response by the autonomous nervous system. The response is generally twofold, including the decrease of heart rate and the reduction of the vascular resistance, both leading to the consequent reduction of blood pressure. Thus the efficacy of this feedback mechanism largely determines the timely and adequate responses to the changes in blood pressure.

Over a long time quantitative assessment of the short-term blood pressure – heart rate feedback regulation efficacy has been limited to the analysis of the baroreflex sensitivity (*BRS*) defined as the measure of the relative change of the pulse interval (in msec) in response to the change in the systolic blood pressure (in mmHg), altogether measured in (msec/mmHg). Historically, the *BRS* was first measured as the increase in the pulse interval to the pharmacologically induced increase in blood pressure that guaranteed the feedback mechanism activation for a certain time period, and the *BRS* was quantified by the linear regression coefficient of pulse intervals on blood pressure over this time period. In the last three decades, several methods to measure *BRS* from simultaneously recorded spontaneous fluctuations of both pulse intervals and blood pressure without applying any external stimuli have been suggested. The most common time-domain approach, often termed as the sequence method, simply focuses on finding time fragments where both pulse intervals and blood pressure either increase or decrease consecutively and monotonously over several heartbeat cycles, sometimes referred to as baroreflex sequences. Next for each of these time fragments a linear regression coefficient of pulse intervals on blood pressure is calculated, representing the local *BRS* estimate [1,2]. To improve the accuracy, averaging over several baroreflex sequences is usually performed, at the cost of lower temporal resolution. In contrast, spectral-based methods do not require selection of certain time fragments, instead focusing on the blood pressure – pulse intervals transfer function analysis in a certain frequency band where the coherence between them exceeds a certain threshold [3–5]. To overcome the common drawback of both methods, in particular their limited performance under non-stationary conditions such as stress tests, modified methods based on first differences analysis have been suggested [6]. Combined with the advances of non-invasive blood pressure measurement techniques, these methods made *BRS* one of the routinely measured parameters both in clinical and ambulatory settings [7].

*BRS* has been reported as a highly informative prognostic marker widely applicable in both ambulatory and clinical investigations. In particular, *BRS* appeared highly predictive of cardiac mortality in post-infarction patients with both reduced ejection fraction [8] as well as preserved left ventricular function [9] including those receiving β-blocker treatment [10] as well as in patients with life-threatening arrhythmias [11], see also [12]. *BRS* impairment has been also reported as an early indicator of autonomic dysfunction and autonomic failure [13]. In earlier studies, *BRS* has been shown to exhibit significant changes under exercise as well as postural and other physical stresses [14–17], while more recent data indicate that these effects are temporary and after a certain adaptation period the baroreflex exhibits a resetting around new absolute blood pressure and pulse interval values [18–20]. However, successful baroreflex resetting has notable exceptions with one such observed recently in diabetes patients with certain complications, particularly with obesity who exhibited significant baroreflex impairment [21].

Of note, *BRS* estimation from either functional test data or spontaneous blood pressure – heart rate oscillation dynamics requires beat-to-beat measurements of blood pressure. When they are not available, a powerful alternative that is competitive in general cardiac prognosis accuracy is provided by the method known as the phase-rectified signal averaging (PRSA) that quantifies the ability of heart rate to either accelerate or decelerate, with the latter being more important for the prognosis, as it is essential for damping critical blood pressure elevations [22]. Simply, physical interpretation of the *PRSA* acceleration/deceleration capacity indices is similar to *BRS*, although the *BRS* is normalized to the ’input’ blood pressure, while *PRSA* is not.

While the *BRS* quantifies explicitly the response of the heart rate to the changes in blood pressure, it does not contain any information whether there was such a response to every significant variation of blood pressure. Time-domain methods simply disregard time intervals without significant changes of heart rate irrelevant to the fact whether there were blood pressure variations, while spectral methods provide characteristics that are averaged over a given frequency band within the entire analysis window. Thus neither of them can guarantee that all blood pressure variations were adequately responded. However, a timely response to the changes in blood pressure is essential for homeostasis, since missing or delayed responses lead to the increased blood pressure variability. In contrast, there is recent evidence that baroreflex activation therapy using implantable devices that stimulates carotid baroreceptors significantly improved the blood pressure control efficacy [23]. Accordingly, additionally to the measurement of the *BRS*, it is also important to quantify its activation in response to blood pressure variations.

In this paper, we suggest a complementary indicator of the blood pressure – heart rate feedback regulation based on their synchronization patterns. We analyzed the synchronous behaviour in both normal subjects and patients with various autonomic regulation dysfunctions under stationary conditions as well as during stress testing. Based on our analysis, we next found time fragments where the standard deviation of the differential instantaneous phases of pulse intervals and blood pressure are below a certain threshold, indicating that they obey synchronous behaviour. In contrast to the sequence method that is based on the linear regression analysis and requires significant changes in the magnitudes of both heart rate and blood pressure, the synchronization analysis is based on the analysis of the instantaneous phases obtained by the Hilbert transform that does not require significant magnitude variations. Thus it also takes into account the hemodynamic variations during those time fragments where the changes of blood pressure and pulse intervals are not monotonous. An additional advantage of the Hilbert transform based phase analysis is that it does not require stationarity of the analysed data series and thus can be straightforwardly applied to stress test records. Furthermore, the estimates are performed in a short time window that potentially allows for the online analysis without any algorithm modification. For an overall quantification of the synchronization behaviour, we finally suggest a synchronization coefficient defined as the fraction of time fragments where the standard deviation of the instantaneous phase difference is below a given threshold within the entire analysis period. In this setting the analysis window may represent a certain phase on stress test such as supine and/or orthostatic phases of a head-up table tilt test. Based on the results of the analysis of synchronization coefficient in different subjects under various conditions, we believe that it could appear an important indicator of the activity of short-term heart rate – blood pressure feedback regulation that is complementary to the *BRS* this way contributing to a deeper understanding and differential diagnostics of autonomic disorders.

Investigation of the physiological signals’ mutual synchronization patterns is widely used in chronobiological studies and is essential for a deeper understanding of the mechanisms related to the influence of various external factors such as geomagnetic field variations, solar cycles, jetlag or night shifts [24–26]. Recent examples include the synchronization analysis of heart rate and respiration during different sleep phases [27–29]. While in most cases relation between rhythms with certain quasi-periodic structure is studied, such as breathing cycles modulating heartbeat cycles, we go beyond that and, while following conceptually similar methodology, modify the synchronization analysis method to suite the blood pressure – pulse interval analysis that both exhibit rather stochastic behavior, as indicated below.

## 2. Materials and Methods

### 2.1 Subjects and experimental protocol

All recordings were obtained at the Almazov National Medical Research Centre in accordance with ethical standards presented in the Declaration of Helsinki. Prior to their inclusion in the study all subjects gave a written informed consent about the protocol. The protocol was reviewed and approved by the Ethics Committee of the Almazov National Medical Research Centre before the beginning of the study.

The main phase of the study included 95 subjects subdivided into 3 groups:

– group 1 included 33 patients aged 39.4±17.5 years old with non-diabetic moderate autonomic dysfunction indicated by previous positive tilt-test responses and orthostatic hypotension characterized by systolic blood pressure (SBP) reduction by at least 20 mmHg and/or by diastolic blood pressure reduction by at least 10 mmHg in the orthostatic compared to supine position;

– group 2 included 34 diabetes mellitus patients aged 52.7±10.9 years old with indications of autonomic neuropathy diagnosed previously by standardized diagnostic tests as indicated in [30,31], with at least two positive test responses required to determine cardiovascular autonomic neuropathy, who received standard combination therapy of blood sugar reducing drugs, including glitazones, gliptins, metformin and sulfonylureas, while no specific therapy of autonomic neuropathy within one month before the study;

– group 3 contained of 28 healthy volunteers aged 41.2±11.1 years old.

All 95 subjects and patients were subjected to head-up tilt table testing (table tilt 60°, duration of orthostatic position up to 30 minutes unless stopped earlier due to syncope response) [32], see also [33] that was performed under identical conditions between 10 am and 1 pm. The recording in the initial supine position and the initial fragment of the orthostatic phase recording (both around 10 min duration) were used in further analysis. Blood pressure recordings were obtained non-invasively with Finometer (TNO, the Netherlands) by the volume-clamp method. From the pulse curve systolic points and pulse intervals between them were extracted, with each pair of data points given by the systolic point and the following pulse interval measurements.

For the initial adjustment of the methods and finding appropriate parameters of the synchronization analysis algorithm that fit to the typical blood pressure and heart rate variability characteristics, another set of 150 stationary records obtained previously from subjects with various autonomic status under supine resting conditions (typical record duration around 10 min) have been used.

### 2.2 Data acqusition and preliminary processing

As the employed instantaneous phase calculation procedure by Hilbert transform is rather sensitive to random errors in measurements, first we applied the adaptive recursive filtering procedure of the original sequence of measurements {*s*_*i*_} (where *s* denotes either *SBP* or *PI*) which aims at the elimination of the anomalous blood pressure and pulse interval measurements. This procedure is based on the analysis of the first differences of the pulse intervals and systolic blood pressure values. The threshold value for the exclusion of outliers is based on the analysis of their empirical distribution functions for the processed data series. The elimination procedure consists of two consecutive steps that are repeated iteratively: (i) marking of the potential outliers as candidates for future elimination and (ii) removing marked outliers. In the first step, one calculates the first differences 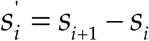 from the initial dataset {*s*_*i*_} (that will be recalculated iteratively each time after a single outlier is eliminated). To mark the *i*-th element of the dataset {*s*_*i*_} as a candidate for being an outlier (“OUT”) three conditions should be met simultaneously:

1. The absolute values of the first differences 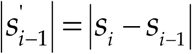 and 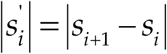 that are adjacent to the measured data {*s*_*i*_} are both either greater or equal than the standard deviation σ{*s*_*i*_} of the first differences.
2. The two consecutive first differences 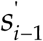 and 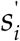 have different signs.
3. The absolute values of the first differences normalized by its standard deviation 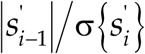 and 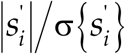 both exceed the critical value σ_crit._ (*N*) determined as a quantile of the Students *t*-distribution for the given dataset size *N*.

Those elements for which all three conditions are met are then marked as “OUT”. Out of the marked data {*s*_*i*_} one with the largest normalized standard deviation is eliminated. After elimination of a single outlier the above procedure is repeated iteratively unless there are no values marked as outliers for the chosen elimination depth. For deeper details on the filtering algorithm, we refer to [34]. The filtering algorithm is also embedded as a separate function in the Matlab source code for *Sync* calculation available as S1 File.

Since the sequence of pulse intervals is non-equidistant due to its inherent variability, we next use the cubic interpolation and resampling with the desired sampling frequency, 5 Hz in our case. Therefore both analysed datasets are now represented by sequences equidistant in time and taken at the same time points.

### 2.3 BRS estimation

To calculate *BRS* from blood pressure and pulse interval recordings during tilt tests, we follow a recently suggested methodology that is particularly suited for dealing with non-stationary data [6]. The first differences of SBP and PI values are taken and the *BRS* is estimated as a linear regression coefficient of ΔPI on ΔSBP in those quadrants where the signs of ΔPI and ΔSBP are identical. To disregard uncertain changes beat-to-beat changes of less than 1 mmHg in SBP or less than 3 ms in PI and more than 20 mmHg in SBP or more than 100 ms in PI are ignored.

### 2.4 PRSA estimation

The *PRSA* index method is based on determining pairs of measurements with either increasing or decreasing pulse interval. For the deceleration capacity estimation from the *PI* measurements one finds all *i* satisfying *PI*_*i*_ < *PI*_*i*+1_ and calculates 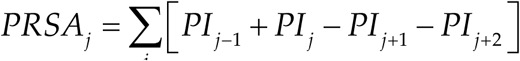 over all selected *i*. Vice versa, to estimate the acceleration capacity one finds all *j* satisfying *PI*_*i*_ > *PI*_*i*+1_ and calculates 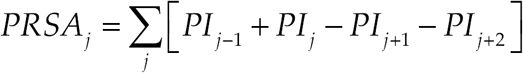. Finally, for the entire analyzed record the *PRSA* index is obtained by simple averaging over all considered *i* and *j*. Accordingly, *PRSA* quantifies the *absolute* acceleration or deceleration capacity of heart rate, while *BRS* characterizes the same capacity *relative* to the input that is determined by the blood pressure change.

### 2.5 The method of PI and BRS phase synchronization measurement

To estimate the mutual synchronization behaviour of the two processes quantitatively we used the method based on the comparison of their phases [35,36]. Instantaneous phase values were determined by Hilbert transform which is widely used in mathematics, physics and signal analysis. For the overall estimation algorithm design, see Fig. 1.

**Figure 1.**
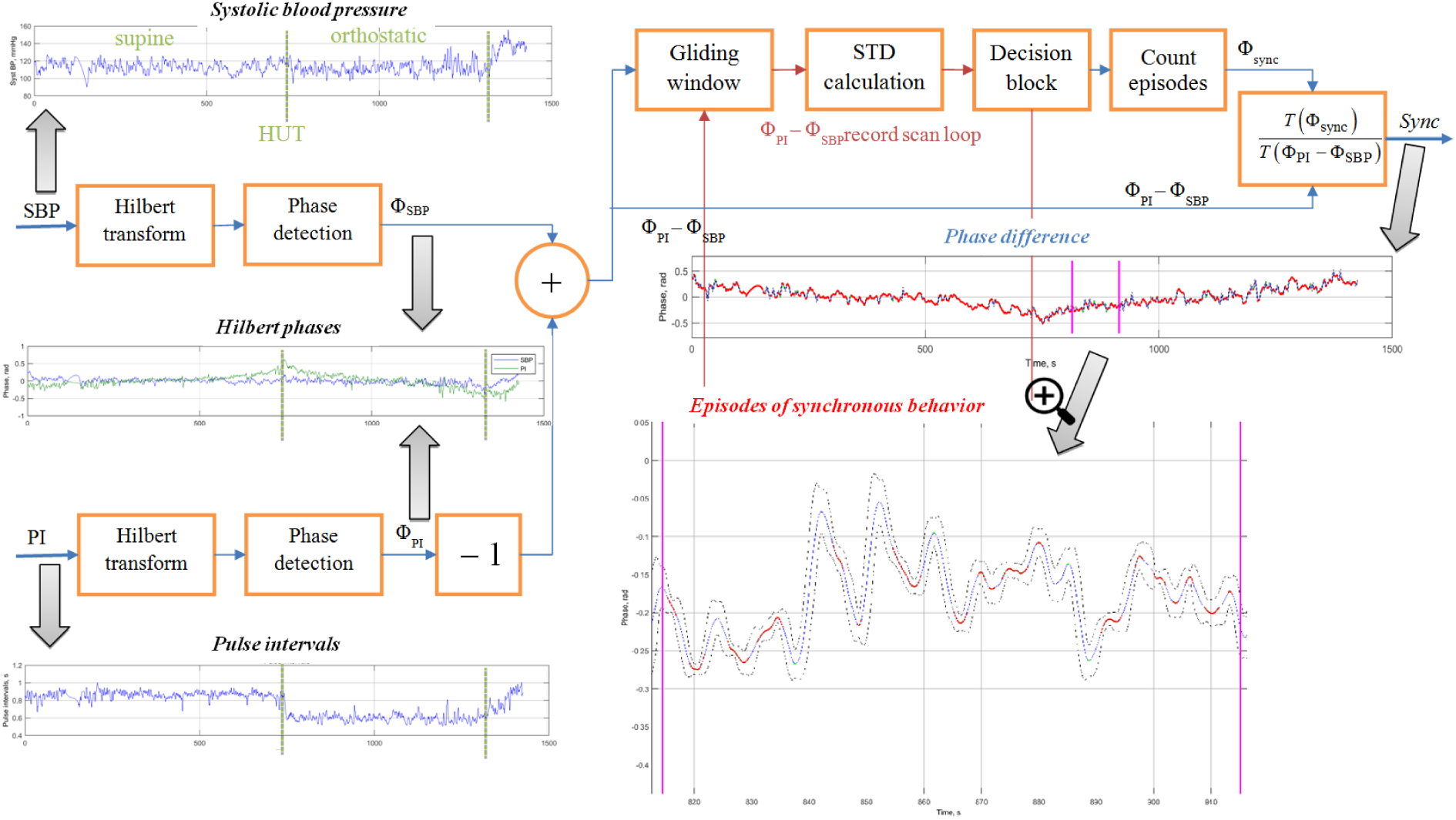
Blood pressure – pulse intervals synchronization analysis algorithm design.

The Hilbert transform produces complex function 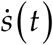 from the original real signal *s*(*t*) (that stands for either *SBP* or *PI*) by adding the imaginary component *s*_⊥_ (*t*), which is defined as

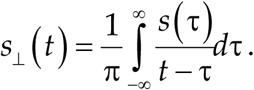

The resulting complex function is known as the complex analytical signal 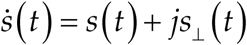.

The real and imaginary parts of the analytical signal allow to determine the envelope *S*(*t*) as the absolute value of the analytic signal that characterizes the laws governing its amplitude modulation, and the phase Φ (*t*) as the argument of the analytical signal that characterizes the laws governing its angular modulation. Accordingly

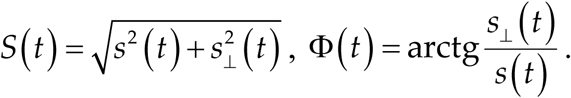

Of note, the signal phase has a clear physical interpretation only in the case of harmonic or narrowband oscillations, while the above formalism is not restricted to these assumptions and thus allows us to calculate phase values for arbitrary data sequences. Next we determine the differences between the phases Φ_PI_ – Φ_SBP_. Following [27, 28] we next applied moving average filtering in a gliding window of size τ and calculated the standard deviation of the phase differences. Consecutive phase points where the standard deviation remains below a given threshold, equal to 2π/δ are treated as belonging to synchronization episodes, once their duration exceeds *T* seconds. This procedure is applied to the entire record in a gliding window while counting episodes of synchronous behavior. As a result, a quantitative measure of the phase synchronization is the synchronization coefficient *Sync* defined as the percentage of the synchronous behaviour episodes duration within the total duration of the analysis window.

Figure 1 exemplifies the entire analysis procedure for a single tilt-test record. After preliminary data preparation both *SBP* and *PI* sequences are subjected to Hilbert transform and phase detection as indicated above. The entire algorithm has been implemented using the Mathworks Matlab software package. A sample of Matlab code that detects synchronization episodes and calculates the synchronization coefficient *Sync* for a given fragment of pairwise blood pressure and pulse intervals measurements is provided as supplementary S1 File. For its immediate testing, a sample Finometer output file with anonymized tilt test record in enclosed as supplementary S2 File.

### 2.6 Adjustment of synchronization analysis algorithm parameters

First we focus on the optimization of synchronization algorithm parameters τ, *T* and δ. The appropriate choice of these values depends on the specific experimental conditions and is not always universal for the given type of data. The parameters may be adjusted to optimize the sensitivity of the algorithm by avoiding the saturation at either very low or very high synchronization coefficients. The gliding window duration τ determines the number of phase points in the window. It is connected with the parameter δ used to calculate the threshold for the standard deviation of the differential instantaneous phase since the standard deviation calculated for a finite data sample depends on the sample size.

To choose the appropriate window size τ, we first focus on the boundary conditions that determine its possible range. One of the common hypotheses of the Mayer waves origin is their baroreflex loop based nature, suggesting that their period of about 10 s corresponds to the full regulation cycle [37]. Accordingly, any internally or externally induced changes in blood pressure are followed by characteristic regulatory oscillations with the Mayer waves period. Thus choosing the gliding window size that is comparable or above this 10 s period will eliminate these short-term regulatory oscillations by averaging. Alternatively, to ensure that the observed variations in both blood pressure and pulse intervals are proper measurements not caused by single faulty measurements and do not appear artifacts of preliminary filtering procedures, we have to guarantee that at least several actual measurements have been performed in each window. Taking into account typical heart rate values, 3 s gliding window will typically result in having 3-4 pulse measurements under resting conditions and even more under stress conditions with increased heart rate. Accordingly, we have to restrict ourselves with τ above 3 s and preferably not longer than half period of Mayer oscillations, that is 5 s. For a better temporal resolution of our analysis, we focus on the lower bound of this range and use 3 s gliding window. Next the threshold for the standard deviation of the phase difference was chosen empirically as 2π/300, *T* = 0.4 s, the latter adjusted empirically to avoid saturation and this way increase the dynamic range of the *Sync* index.

The graphic insets in Fig. 1 exemplify the analysis results for a single tilt-test record. In addition to preprocessed data sequences and their Hilbert phases, the figure displays the differential phase as well as the results of its standard deviation analysis in a gliding window (in the lower right panel). The dashed lines denote the 2π/δ threshold for the standard deviation of the phase difference. The bold solid curve denotes Φ_PI_ – Φ_SBP_ and highlights (in red) the synchronization episodes where the standard deviation of the phase difference remains below 2π/δ threshold. Finally, the fraction of such episodes within the total record duration determines the synchronization coefficient *Sync*.

The figure shows that, while for the entire analysed fragment the synchronization coefficient *Sync* is somewhat around 50%, it exhibits considerable variations along the record. While the first part of the record is characterized by prolonged episodes of synchronous behaviour interrupted by few short-term asynchronous fragments, the second part of the fragment exhibits more frequent asynchronous behaviour episodes interrupted by rather few coupling patterns. While the particular reasons for each onset and breakdown of this coupling can hardly be determined, there is a clear discrepancy between the first and the second part of the record in terms of their synchronization patterns. Such changes in the synchronization behaviour of blood pressure and pulse intervals could be triggered by some physical or mental stress, change in the body position, and so on. Accordingly, reactions to various stress patterns that are imposed during functional tests can be studied in terms of the changes in the blood pressure – pulse intervals synchronization coefficient *Sync*. This requires the recordings being analysed during different test phases. For example, for the head-up tilt (HUT) table testing the supine and the orthostatic test phases could be analysed separately, this way allowing to evaluate how the synchronization patterns changes in response to the orthostatic stress. An additional advantage of the proposed methodology is that the evaluation of the degree of phase synchronization between PI and SBP may be useful in the study of regulatory functions in the human body during various functional tests, since it does not require data stationarity.

While the *BRS* value characterizes the *intensity* of the heart rate reaction to the blood pressure changes during observed feedback responses, the *Sync* value likely indicates how often such responses are *activated* in the first place. Accordingly, *low intensity* of the reaction characterized by low *BRS* that should result in higher than normal blood pressure variability, theoretically could be at least partially compensated by its *more frequent activation* characterized by higher *Sync*.

### 2.7 Statistical analysis

Since our preliminary studies [35,36,38] indicated that the synchronization coefficient is not normally distributed, we used the methods of non-parametric statistics to process our results. To quantify the statistical significance of our results, we next applied the non-parametric Mann-Whitney U-test and the non-parametric Kolmogorov-Smirnov test for independent samples [39]. To calculate the correlations between the synchronization coefficient, the standard deviations of pulse intervals and systolic blood pressure and the baroreflex sensitivity we used the non-parametric Spearman’s correlation coefficients. For those cases where the dependence appeared significant in the first place (*p* < 0.01), we next applied linear regression analysis with an iterative outlier elimination procedure following a standard methodology [40], used also previously in [41]. All statistical analysis was performed using the IBM SPSS Statistics software package.

## 3. Results

### 3.1 Relation between BRS, Sync, blood pressure and heart rate variability under stationary conditions

Next using the methodology proposed above we examined both *BRS* and *Sync* coefficient in subjects with various autonomic status under stationary conditions. Figure 2 shows the normalized histograms that provide the probability density function (PDF) estimates (A-C) and the cumulative distribution functions (CDF) (D-F) of the synchronization coefficient *Sync* (*med* = 83; *iqr*= 19), the average baroreflex sensitivity *BRS* (*med* = 5.25; *iqr* = 4.29) as well as the product of *Sync* and *BRS* (*med* = 405.25; *iqr* = 363.02) under stationary conditions, where *med* are the medians and *iqr* are the interquartile ranges, respectively. The figure clearly shows that all empirical distributions are strongly non-Gaussian. The analysis of the *Sync* distribution indicates that the values between 80 and 95 are the most probable, indicating a large fraction of synchronous episodes in the studied records.

**Figure 2.**
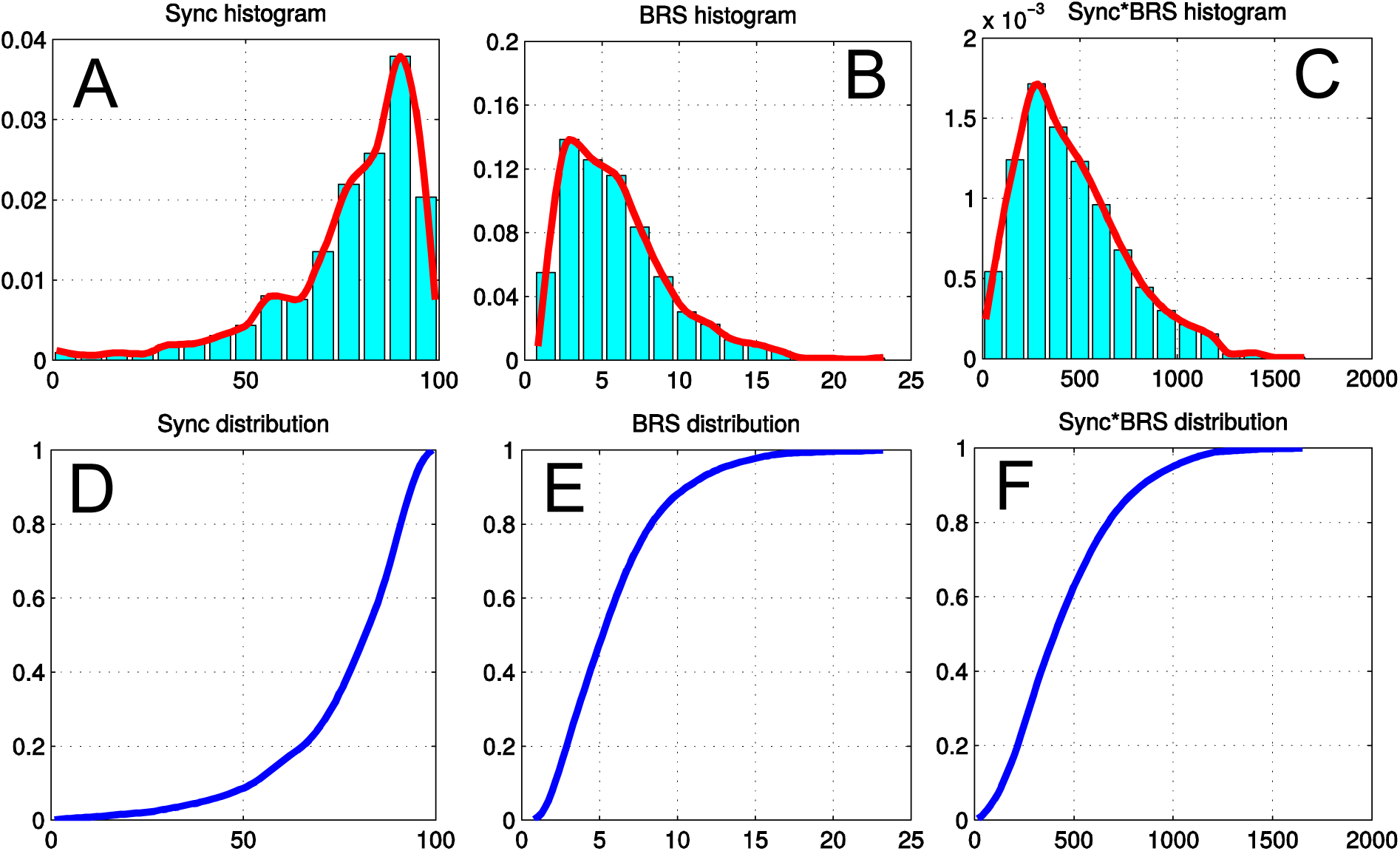
(A) The synchronization coefficient histogram, (B) the baroreflex sensitivity histogram, (C) the product of *Sync* and *BRS* histogram, (D) the synchronization coefficient CDF, (E) the baroreflex sensitivity CDF, (F) the product of *Sync* and *BRS* CDF.

Figure 3 shows that, while BRS is positively correlated with the standard deviation of the pulse intervals (the non-parametric Spearman’s correlation coefficient *C* = 0.66, significance level *p* < 0.001), in marked contrast *Sync* exhibits pronounced negative correlations (*C* = −0.65, *p* < 0.001). This may indicate that under reduced *BRS* conditions when the heart rate does not change sufficiently in response to blood pressure variations leading to lower pulse interval variance, higher *Sync* indicates more frequent activation of the feedback responses to partially compensate the reduced *BRS*. Finally, the *BRS*\**Sync* product also increases with increasing *PI* standard deviation. Accordingly, the *BRS*\**Sync* product seems a proper indicator of the overall contribution of the feedback mechanisms to maintain haemostatic BP values that takes into account both the sensitivity of the reflexes as well as their total activity time.

**Figure 3.**
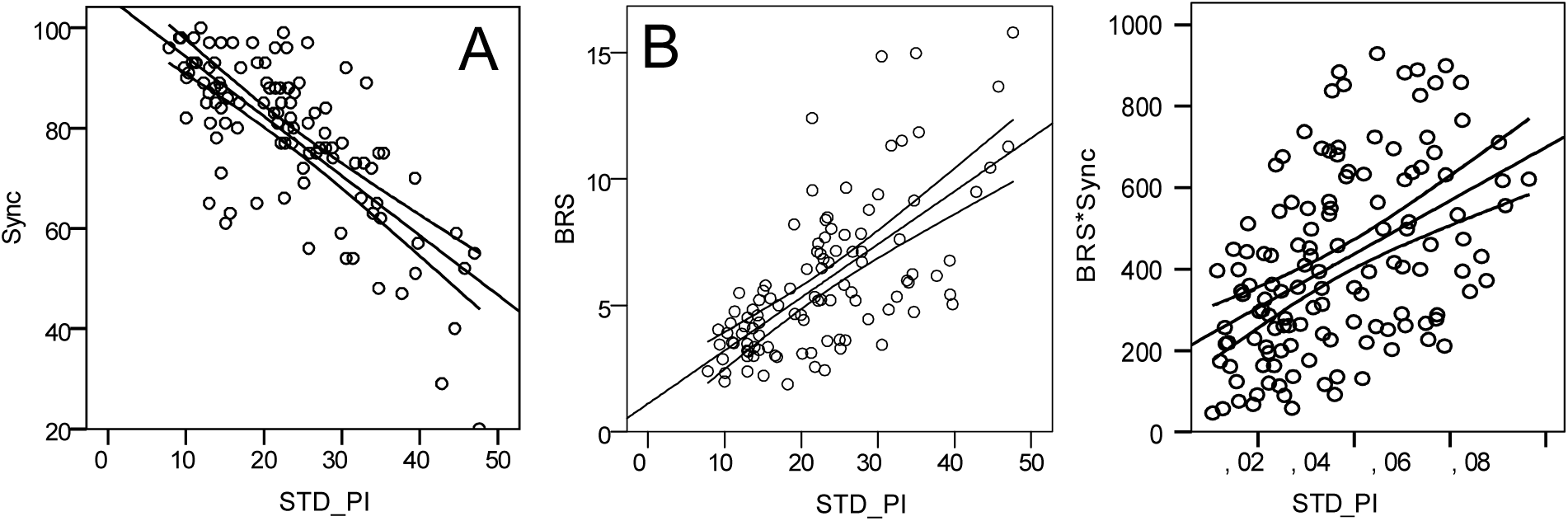
Linear regressions between (A) *Sync* and *PI* variance, (B) *BRS* and *PI* variance, (C) logarithms of *BRS*\**Sync* product and of SBP standard deviation. Relevant regression lines and their 95% confidence intervals are shown in each panel.

### 3.3 Blood pressure and heart rate dynamics during tilt test

Next we have analysed the data for the three groups of patients, in particular 33 patients with non-diabetic moderate autonomic dysfunction (group 1), 34 diabetic patients (group 2) and 28 healthy volunteers (group 3).

Figure 4 shows that in both patient groups (1 and 2) SBP during tilt test exhibited considerably more pronounced variations than in the control group, confirming limitations in their BP control. Although the typical (median) SBP variability appeared rather comparable in groups 1 and 2 (see Fig. 4A), the diabetes patients were characterized by significantly lower pulse interval variability (see Fig. 4B). In addition, insufficient pulse interval regulation is also indicated by reduced PRSA index (see Fig. 4C).

**Figure 4.**
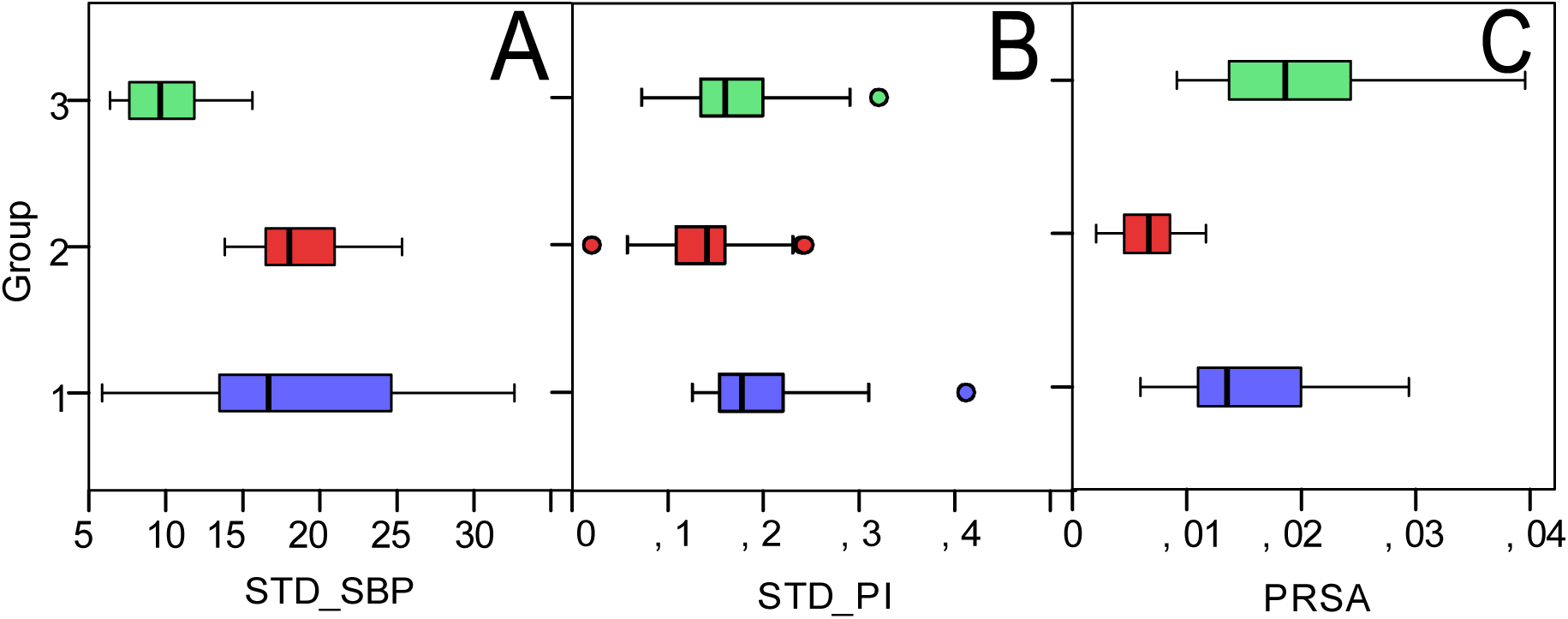
Boxplots of (A) *STD_SBP*, (B) *STD_PI* and (C) *PRSA* for the entire tilt test records. Group 1 – patients with non-diabetic moderate autonomic dysfunction, group 2 – diabetic patients with confirmed autonomic neuropathy, group 3 – healthy volunteers.

Analysis of the entire tilt test records revealed that, as expected, the *BRS* differs significantly in healthy individuals and in patient groups as well as between diabetic patients and the broader patient group with moderate autonomic disorders (see Fig. 5). Statistical characteristics of the estimated distribution parameters for all studied quantities (mean, standard deviation, median and interquartile range) obtained for the entire tilt test records for all studied groups are summarized in the Table A1 in the Appendix.

**Figure 5.**
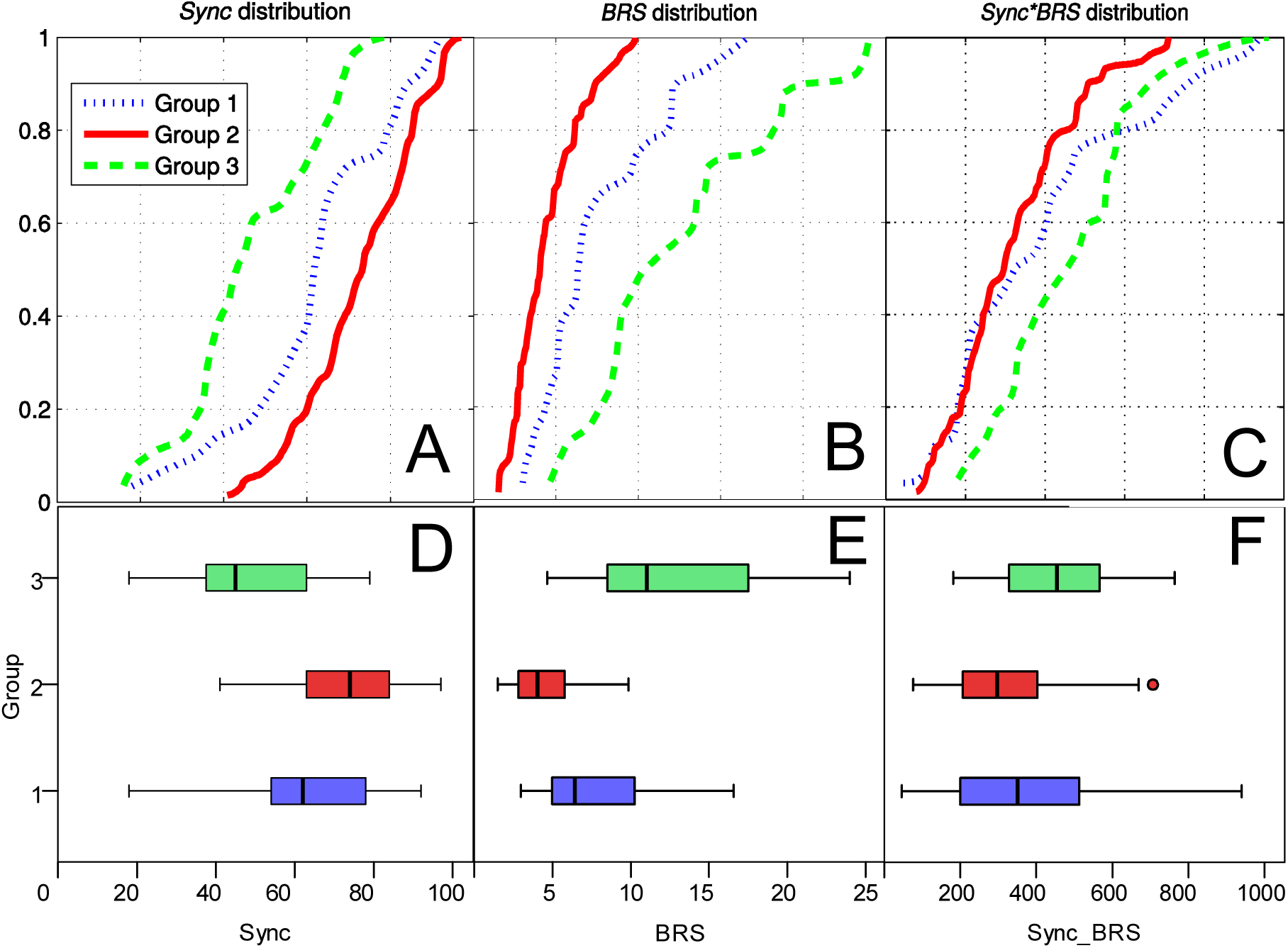
CDFs and boxplots of (A, D) *Sync*, (B, E) *BRS* and (C, F) *BRS*\**Sync* product (upper and lower panels, respectively), for the entire tilt test records. Group 1 included patients with non-diabetic moderate autonomic disorders, group 2 included diabetic patients with confirmed autonomic neuropathy, while group 3 included healthy volunteers.

Both the non-parametric Mann-Whitney U-test and the Kolmogorov-Smirnov test for independent samples confirmed significant differences between the studied parameters in various groups. We found that the synchronization coefficient *Sync* is significantly higher and the baroreflex sensitivity *BRS* is significantly lower in the group 2 (diabetic patients) in comparison with other groups of patients for the entire test records (significance level *p*_U_ < 0.001 based on the Mann-Whitney U-test; significance level *p*_KS_ < 0.01 based on the Kolmogorov-Smirnov test). The synchronization coefficient *Sync* is significantly lower and the baroreflex sensitivity *BRS* is significantly higher in group 3 (healthy volunteers) than in groups containing patients with different autonomic disorders (*p*_U_ < 0.001; *p*_KS_ < 0.01). The *BRS*\**Sync* product is also significantly higher in group 3 than in the group 2 (*p*_U_ < 0.005; *p*_KS_ < 0.05), while no significant differences between groups 1 and 3 could be observed.

Figures 6 illustrates linear regressions with 95% confidence interval between the synchronization coefficient *Sync* and the average baroreflex sensitivity *BRS.* Results for the diabetes patients are shown in Fig. 6A while results for patients from other groups and for healthy subjects are shown in Fig.6B. Similar results are shown for the *PRSA*, an index counterpart to BRS while not normalized to the ’input’ blood pressure, in Figs. 6C and 6D.

**Figure 6.**
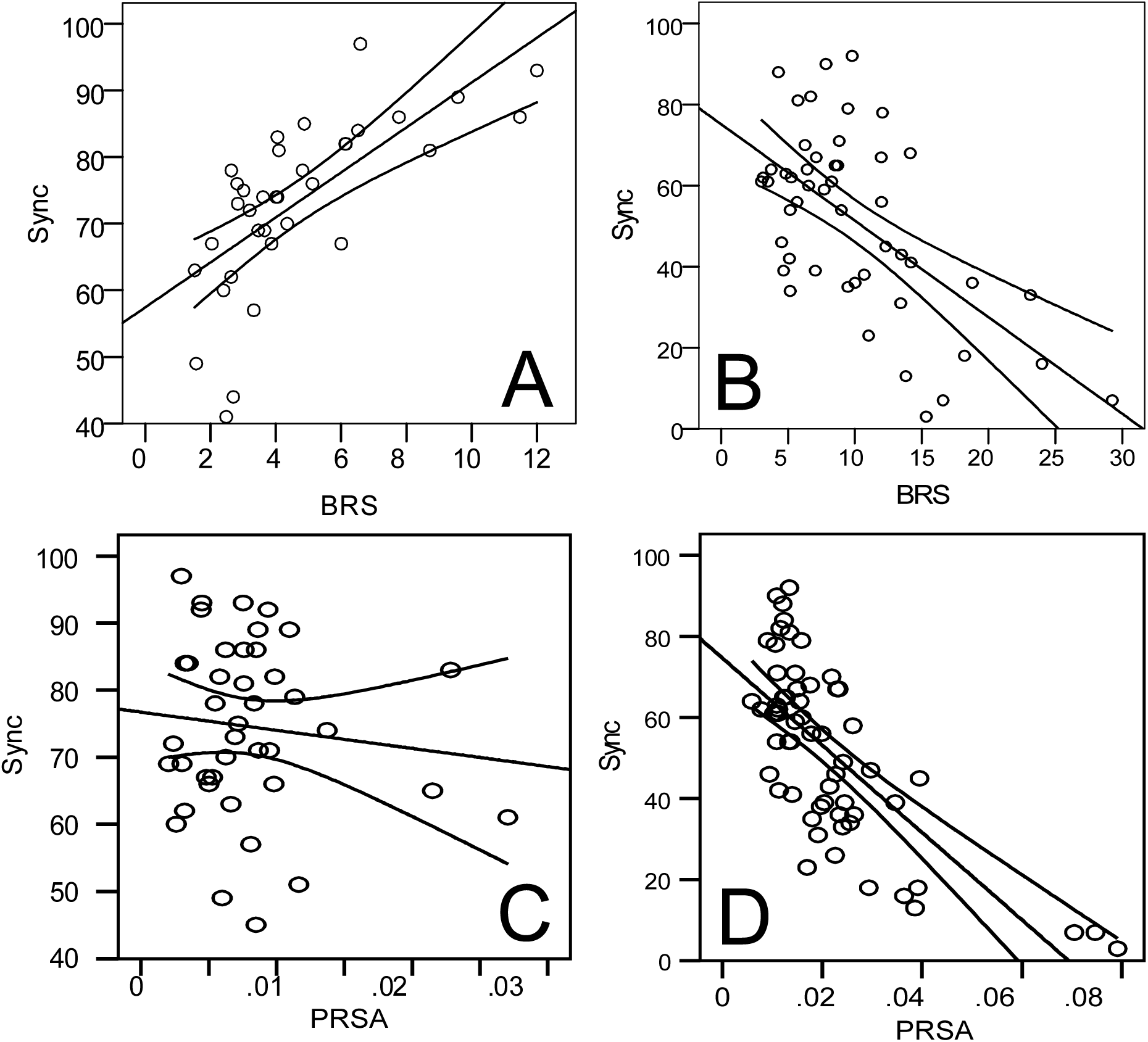
(A) Linear regression between *Sync* and *BRS* in diabetic patients, (B) linear regression between *Sync* and *BRS* in patients with non-diabetic moderate autonomic dysfunction and healthy volunteers, (C) linear regression between *Sync* and *PRSA* in diabetic patients, (D) linear regression between *Sync* and *PRSA* in patients with non-diabetic moderate autonomic dysfunction and healthy volunteers.

We found significant positive correlation between the synchronization coefficient *Sync* and the average baroreflex sensitivity *BRS* in group 2 diabetic patients for the entire records (*C* = 0.82, significance level *p* < 0.001). In contrast, in other groups, this relation is reversed, indicated by a negative correlation coefficient with significantly lower absolute value. In healthy volunteers and patients with moderate autonomic disorders studied together *C* = –0.47 (*p* < 0.001). Again, like in the stationary data analysis, we tend to attribute this effect to the partial compensation of the reduced *BRS* by a more frequent activation of the feedback responses. In contrast, in diabetes patients this compensation mechanism is likely disrupted leading to further deficiencies in blood pressure control. Qualitatively similar results have been obtained for the *PRSA*, an index counterpart to BRS while not normalized to the ’input’ blood pressure, in both healthy subjects and patients with moderate autonomic disorders, while in diabetes patients no significant correlation between *PRSA* and *Sync* could be observed.

### 3.4 Blood pressure and heart rate dynamics during supine pre-tilt phase

Comparison of the studied indices for the supine phase of the tilt test indicates that both BRS and *Sync* also differ significantly between healthy individuals and two patient groups as well as between diabetic patients and non-diabetic patients. Boxplots and distribution plots in Fig. 7 illustrate the differences in the distribution characteristics of the synchronization coefficient, baroreflex sensitivity and the *BRS*\**Sync* product, estimated for the supine phase of the tilt test for the selected patient groups. Statistical characteristics of the corresponding estimates (mean, standard deviation, median and interquartile range) are summarized in Table 1 in the Appendix.

**Figure 7.**
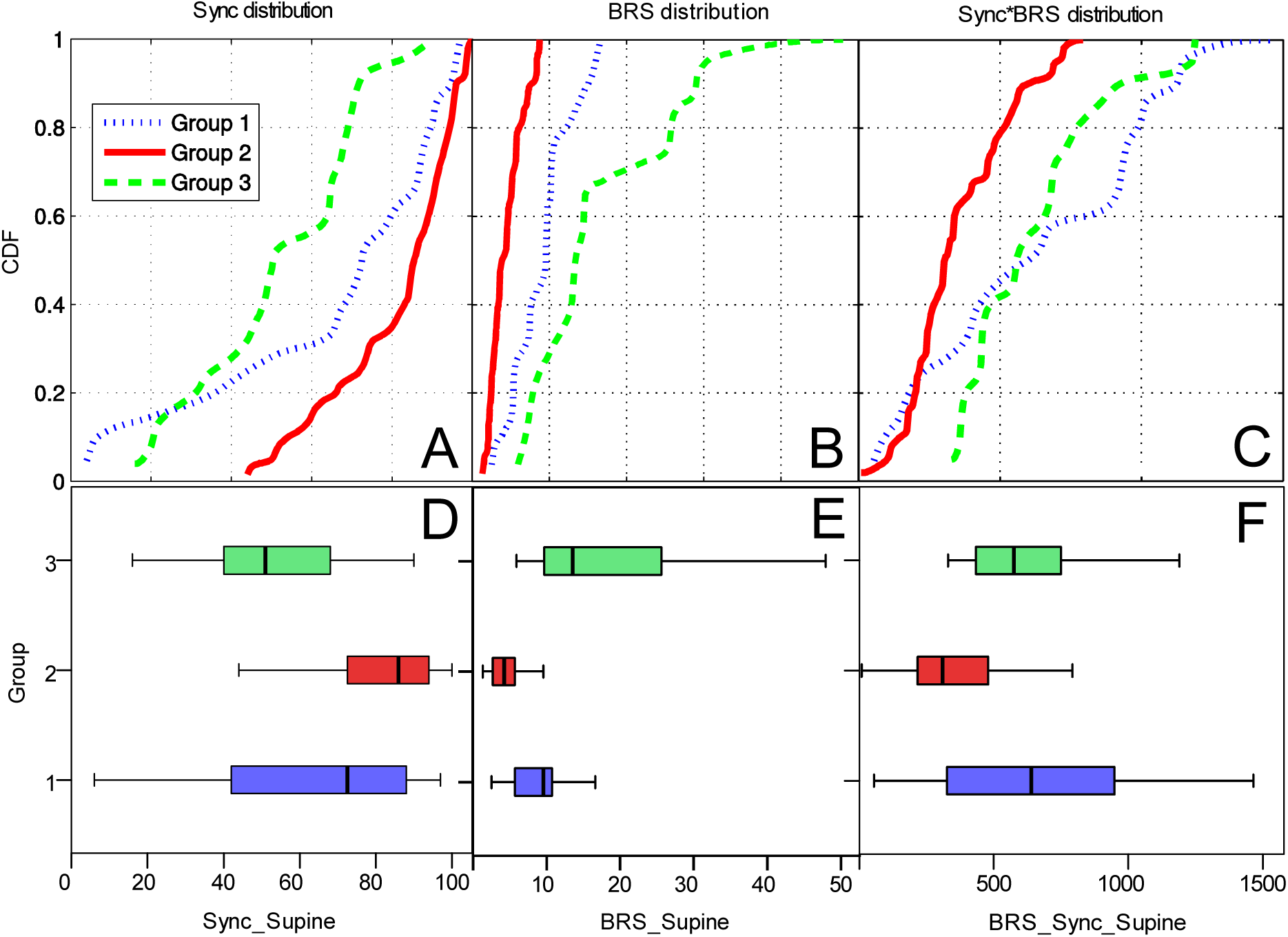
The synchronization coefficient CDFs (A) and boxplots (D), the baroreflex sensitivity CDFs (B) and boxplots (E), the *BRS*\**Sync* product CDFs (C) and boxplots (F), estimated for the supine phase of the tilt test for the selected patient groups. Group 1 included patients with non-diabetic moderate autonomic disorders, group 2 included diabetic patients with confirmed autonomic neuropathy, while group 3 included healthy volunteers.

The non-parametric Mann-Whitney U-test and the Kolmogorov-Smirnov test for independent samples confirmed significant differences between the studied parameters in patient groups. The baroreflex sensitivity *BRS* is significantly lower in the group 2 (diabetic patients) in comparison with other groups of patients for the supine phase of the tilt test (*p*_U_ < 0.001; *p*_KS_ < 0.001). The synchronization coefficient *Sync* is significantly higher in the group 2 then in the group 3 (*p*_U_ < 0.001; *p*_KS_ < 0.001). The synchronization coefficient *Sync* is significantly lower in the group 3 (healthy volunteers) compared with other groups of patients (*p*_U_ < 0.001; *p*_KS_ < 0.01) and the baroreflex sensitivity *BRS* is significantly higher (*p*_U_ < 0.001; *p*_KS_ < 0.02). The product of *Sync* and *BRS* is significantly higher in the group 3 then in group 2 (*p*_U_ < 0.001; *p*_KS_ < 0.001).

We found moderate significant negative correlations between the synchronization coefficient *Sync* and the standard deviation of SBP in diabetic patients (*C* = –0.43, *p* < 0.001) as well as between the *BRS*\**Sync* product and the standard deviation of SBP (*C* = –0.38, *p* < 0.05) in other groups.

### 3.5 Blood pressure and heart rate dynamics during post-tilt orthostatic phase

Boxplots and CDFs in Fig. 8 illustrate the differences in the distribution characteristics of the synchronization coefficient and baroreflex sensitivity between the diabetic patients and other groups, estimated for the orthostatic phase of the tilt test. Statistical characteristics of the corresponding estimates (mean, standard deviation, median and interquartile range) are summarized in Table A1 in the Appendix.

**Figure 8.**
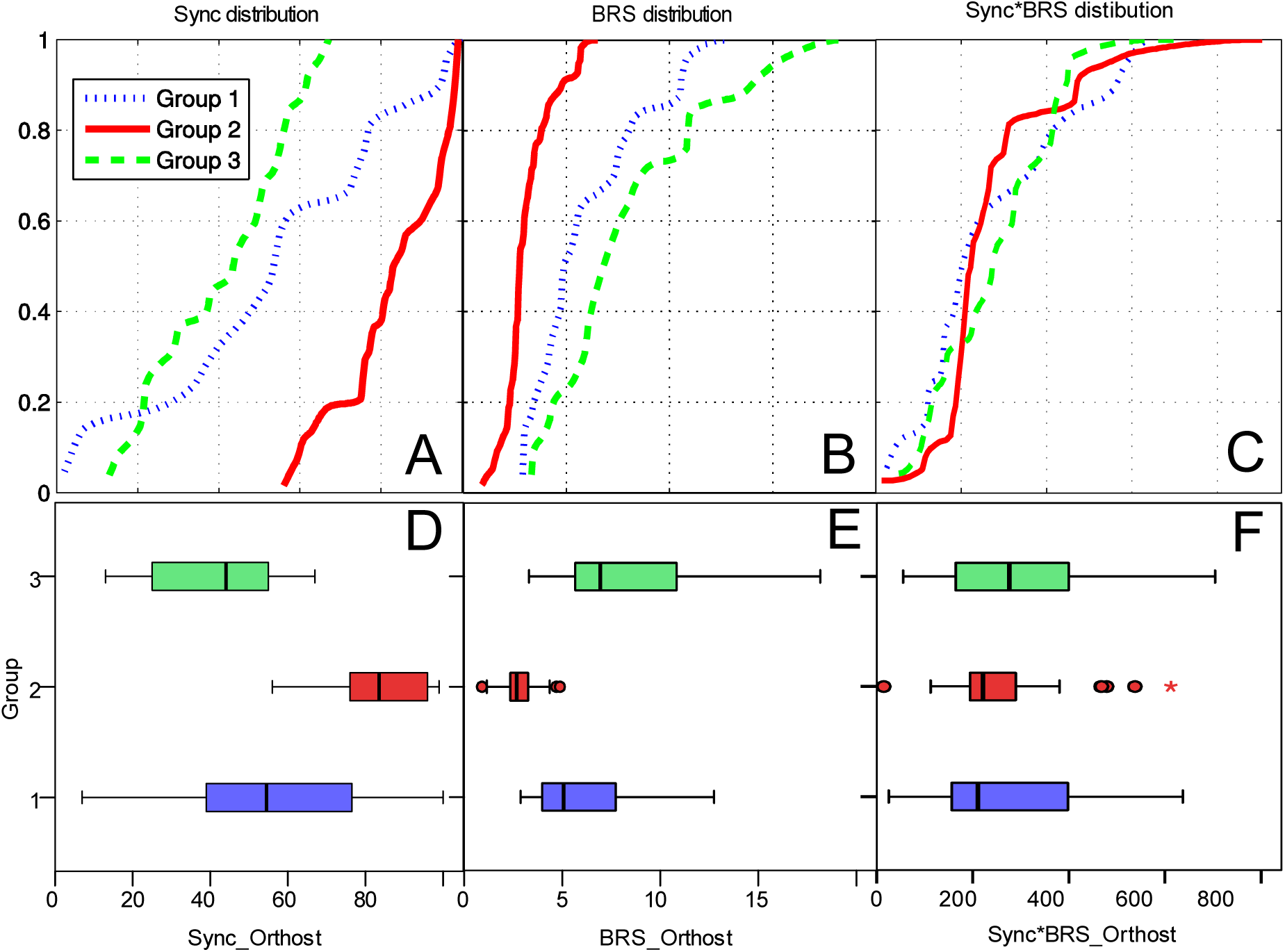
The synchronization coefficient CDFs (A) and boxplots (D), the baroreflex sensitivity CDFs (B) and boxplots (E), (c) the *BRS*\**Sync* product CDFs (C) and boxplots (F), estimated for the supine phase of the tilt test for the selected patient groups estimated for the orthostatic phase of the tilt test for the selected patient groups. Group 1 included patients with non-diabetic moderate autonomic disorders, group 2 included diabetic patients with confirmed autonomic neuropathy, while group 3 included healthy volunteers.

The synchronization coefficient *Sync* is significantly higher and *BRS* is significantly lower in the diabetic patients then in healthy volunteers and in patients with non-diabetic moderate autonomic dysfunction (*p*_U_ < 0.001; *p*_KS_ < 0.001).

We found both pronounced and significant negative correlation between the *BRS*\**Sync* product and the standard deviation of the systolic blood pressure *STD_SBP* (*C* = –0.78, *p* < 0.001), moderate negative correlation between *BRS* and *STD_SBP* (*C* = –0.52, *p* < 0.001) in healthy volunteers as well as moderate negative correlation between *Sync* and *STD_SBP* (*C* = –0.44, *p* < 0.001) in all patient groups.

### 3.6 Comparison between supine and orthostatic tilt test phases

Patients with non-diabetic moderate autonomic dysfunction, diabetic patients and healthy volunteers demonstrate different reactions to the tilt test. We found that the baroreflex sensitivity *BRS* and, as a consequence, also the *BRS*\**Sync* product significantly decreases during the transition to the orthostatic phase of the tilt test (*p*_U_ < 0.001 and *p*_KS_ < 0.001 for diabetic patients; *p*_U_ < 0.01 and *p*_KS_ < 0.05 for the patients with non-diabetic moderate autonomic dysfunction; *p*_U_ < 0.001 and *p*_KS_ < 0.002 for healthy volunteers).

Only in healthy volunteers the standard deviation of the systolic blood pressure *STD_SBP* significantly increases during the transition to the orthostatic phase (*p*_U_ < 0.001; *p*_KS_ < 0.02). Moreover in this group the synchronization coefficient *Sync* is significantly lower in the orthostatic phase of the tilt test than in the supine phase (*p*_U_ < 0.05). Therefore in healthy people the *BRS*\**Sync* product is lower in the orthostatic phase not only because of the lower *BRS*, but also due to the lower *Sync* values.

## 4. Discussion

While key indicators of the cardiovascular feedback regulation are characterized by well-known measures such as *BRS*, some details could not be revealed by using this single indicator. In particular, *BRS* exhibited only moderate reduction in patients with non-diabetic moderate autonomic disorders (group 1), while was reduced considerably in diabetic patients with autonomic neuropathy (group 2). According to these data, one could expect higher blood pressure variability in group 2, as their autonomic nervous system is less sensitive to changes in blood pressure, and their feedback response to blood pressure variations is weaker than in group 1. Surprisingly, these groups were characterized by comparable blood pressure variability with only slightly lower pulse intervals variability in group 2 (see Figs. 4A and 4B), indicating that there might be another reason for that.

That reason could be revealed when considering the introduced synchronization coefficient *Sync* that indicates the (normalized) total duration of the time fragments when mutually synchronized behavior between blood pressure and pulse intervals can be observed. Our results indicate that in all studied patient groups under normal conditions *Sync* is reciprocal to *BRS* indicating that in patients with reduced *BRS* the feedback mechanisms have to be activated more frequently this way trying to compensate lower sensitivity and intensity of feedback mechanisms by their more frequent activation. This compensation seems to be rather a universal phenomenon, as the reciprocal character of *Sync* and *BRS* can be clearly observed in the comparison between them in different studied groups in both supine and orthostatic positions (see boxplots in Figs. 5, 7 and 8). The reciprocal relations between *Sync* and *BRS* are further supported by the regression analysis in groups 1 and 3 (see Fig. 6B). In contrast, in the diabetes patients (group 2) just the opposite can be observed, indicating a breakdown of this compensation in diabetes patients with autonomic neuropathy under certain conditions.

Another way to support the compensation of lower *BRS* by the more frequent activity of the corresponding feedback mechanisms is by considering the *BRS*\**Sync* product. This derived quantity could be interpreted through a simple mechanistic analogy. The *BRS* characterizes the intensity of the baroreflex response. Physiological system with higher *BRS* responds more rapidly and intensively to blood pressure variations and thus requires less time to compensate them. Thus the *BRS* could be considered as a certain equivalent of the response power. In contrast, *Sync* serves as an analog of the total time this power has been applied. In these terms the *BRS*\**Sync* product would be analogous to the amount of work that has been performed by the regulatory system to compensate the blood pressure variations.

The fact that both *BRS* and *Sync* are essential for appropriate short-term blood pressure regulation is strongly supported by their negative significant correlations with blood pressure variability. These correlations are most pronounced in orthostatic position when both timely activation and efficacy of the baroreflex feedback is essential for the short-term blood pressure regulation. It is therefore not surprising that the *BRS*\**Sync* product exhibits most pronounced negative correlations with blood pressure variability. Through the prism of the simple mechanistic analogy mentioned above, low blood pressure oscillations in the orthostatic position are associated with the largest amount of work performed by the regulatory feedback mechanisms, given by the product of their power characterized by *BRS* and their activity time characterized by *Sync*.

Under moderate autonomic dysfunction characterized by reduced *BRS*, more frequent responses to the blood pressure variations are characterized by increased *Sync*, this way keeping their product, while moderately reduced compared to healthy subjects, nevertheless sufficiently high indicating altogether more or less adequate responses to orthostatic stress (see Fig. 5F and Fig. 7F). In contrast, in diabetes patients in orthostatic position we observed a very broad distribution of *BRS*\**Sync* product with outliers at both ends (see Fig. 8F). This might indicate that, while in some patients there is sufficient compensation bringing them in the same range of *BRS*\**Sync* product with normal subjects and non-diabetic patients, other diabetic patients from the same group were characterized by both *BRS* and *Sync* reduced likely indicating that no relevant compensation did occur.

This result is generally in line with recent studies of baroreflex control in diabetes patients that indicated significant dependence of the orthostatic baroreflex performance on concomitant conditions such as obesity [21]. Moreover, in the same study a significantly higher number of baroreflex sequences indicating more frequent activation of the baroreflex loop could be observed in diabetes patients with concomitant obesity compared to control subjects, although differences between diabetes patients and weight-matched control subjects appeared insignificant. Since we have observed similar effect in our study by using the suggested *Sync* indicator, we believe concomitant conditions such as obesity are likely playing a key role in the impairment of baroreflex control not only in terms of its low sensitivity, but also in terms of its timely activation in response to blood pressure variations.

## 5. Conclusion

To summarize, we have analysed blood pressure – heart rate synchronization dynamics in both normal subjects and patients with various autonomic disorders under both stationary and orthostatic stress test conditions. To find the episodes of synchronous behaviour, we suggested the methodology based on the instantaneous phase analysis assessed by Hilbert transform. Although a similar approach has been recently used in the studies of heart rate and respiration mutual behaviour that both exhibit quasi-periodic patterns [27,28], we have adjusted this methodology to best characterize the heart rate and blood pressure mutual dynamics which exhibit rather stochastic behavior. We determined the synchronization coefficient *Sync* as the percentage of time where the standard deviation of difference between instantaneous phases of blood pressure and pulse intervals is below a certain threshold. While the widely adopted *BRS* index characterizes the *intensity* of the heart rate response to the blood pressure changes during observed feedback responses, the *Sync* value indicates how often blood pressure and pulse intervals are synchronized, indicating that correspoding feedback responses are *activated* in the first place. We found that in both healthy subjects and patients with moderate autonomic dysfunction *BRS* and *Sync* are typically reciprocal suggesting that *low BRS* is typically compensated by the *more frequent activation* of the baroreflex loop indicated by higher *Sync*. In contrast, in diabetes patients with autonomic neuropathy *BRS* and *Sync* are positively correlated likely indicating the breakdown of this compensation in some of the diabetic patients with certain concomitant conditions. Therefore we suggest that *Sync* could be used as an additional indicator of blood pressure – heart rate feedback regulation activity that is complementary to the widely used baroreflex sensitivity (*BRS*).

## Appendix

**Table A1.** All the data are presented in the format of the median (interquartile range) that notes in the table as med (iqr) and also as the mean value ± standard deviation that notes as mean ± std. Grey cells mark all cases with significantly different values for diabetic patients and other groups detected by non-parametric Mann-Whitney U-test (significance level no worse than *p* < 0.05) [39]. Bold font indicates all cases when parameters significantly changes during the transition from the supine phase to the orthostatic phase of the tilt test (significance level *p* < 0.05 based on the Mann-Whitney U-test).

| Measured characteristics |  | STD_SBP |  | BRS |  | Sync |  | BRS*Sync |  |
| --- | --- | --- | --- | --- | --- | --- | --- | --- | --- |
| Patient groups | | med<br>(iqr) | mean<br>± std | med<br>(iqr) | mean<br>± std | med<br>(iqr) | mean<br>± std | med<br>(iqr) | mean<br>± std |
| Patients with non-diabetic moderate autonomic dysfunction (group 1) | the entire record | 16.65<br>(11.6) | 19.08±<br>10.12 | 6.42<br>(5.26) | 7.67±<br>3.85 | 62<br>(24) | 62.34±<br>18.39 | 350<br>(292) | 394±<br>248 |
| | the supine phase of the tilt test | 7.83<br>(4.13) | 7.81±<br>2.72 | **9.59**<br>**(4.65)** | **9.23±**<br>**3.94** | 73<br>(37) | 66.33±<br>29.34 | **639**<br>**(623)** | **644±**<br>**406** |
| | the ortho-static phase of the tilt test | 7.69<br>(4.71) | 7 ±<br>3.11 | 5.16<br>(4.12) | 7.26±<br>5.18 | 54<br>(39) | 54.6 ±<br>29.31 | 210<br>(260) | 299±<br>254 |
| Diabetic patients with autonomic neuropathy (group 2) | the entire record | 18.01<br>(5.98) | 18.93±<br>6.71 | 4.05<br>(2.84) | 4.48±<br>2.11 | 74<br>(21) | 73.05±<br>14 | 298<br>(199) | 319±<br>157 |
| | the supine phase of the tilt test | 5.58<br>(3.08) | 6.47±<br>3.02 | **4.07**<br>**(2.98)** | **4.44 ±**<br>**2.1** | 86<br>(21) | 81.52±<br>15.55 | **309**<br>**(259)** | **352±**<br>**183** |
| | the ortho-static phase of the tilt test | 6.15<br>(1.84) | 6.05±<br>1.84 | 2.73<br>(1.03) | 3.07±<br>1.19 | 83.5<br>(19.7) | 83.36±<br>13.41 | 245<br>(272) | 372±<br>349 |
| Healthy volunteers (group 3) | the entire record | 9.65<br>(4.79) | 10.32±<br>4.04 | 11.02<br>(8.99) | 12.62±<br>5.85 | 44<br>(26) | 47.32±<br>16.8 | 469<br>(247) | 471±<br>190 |
|  | the supine phase of the tilt test | **4.80**<br>**(2.67)** | **4.87 ±**<br>**2.67** | **13.6**<br>**(15.9)** | **17±**<br>**9.9** | **51**<br>**(28)** | **52.8 ±**<br>**20.0** | **575**<br>**(305)** | **628±**<br>**254** |
| | the ortho-static phase of the tilt test | 7<br>(3.97) | 7.32±<br>2.34 | 6.9<br>(5.2) | 8.1±<br>4.0 | 44<br>(30) | 41.3±<br>15.3 | 275<br>(234) | 288±<br>150 |

## References

[1] Bertinieri G, Di Rienzo M, Cavallazzi A, Ferrari AU, Pedotti A and Mancia G A new approach to analysis of arterial baroreflex J Hypertension 1985; 3 suppl. 3: 79–81

[2] Malberg H, Wessel N, Hasart A, Osterziel KJ and Voss A Advanced analysis of spontaneous baroreflex sensitivity, blood pressure and heart rate variability in patients with dilated cardiomyopathy Clin Sci 2002; 102 : 465–473

[3] De Boer RW, Karemaker JM and Strackee J Relations between short-term blood pressure fluctuations and heart rate variability in resting subjects: A spectral analysis approach Med Biol Eng Comp 1985; 23 : 352–364

[4] Robbe HW, Mulder LJ, Ruddel H, Langewitz WA, Veldman JB and Mulder G Assessment of Baroreceptor Reflex Sensitivity by Means of Spectral Analysis Hypertension 1987; 10 :538–543

[5] Cerutti S, Baselli G, Civardi S, Furlan R, Lombardi F, Malliani A, Merri M and Pagani M Spectral analysis of heart rate and blood pressure variability signals for physiological and clinical purposes Proc Comp Card 1987; pp. 435–438

[6] Bogachev MI, Mamontov OV, Konradi AO, Uljanitski YD, Kantelhardt JW and Schlyakhto EV Analysis of blood pressure–heart rate feedback regulation under non-stationary conditions: beyond baroreflex sensitivity Physiol Meas 2009; 630 : 631–645

[7] Parati G, Omboni S, Fratolla A, Di Rienzo M, Zanchetti A and Mancia G Dynamic evaluation of baroreflex in ambulant subjects Blood Pressure and Heart Rate Variablity, edited by M. Di Rienzo et al. Amsterdam: IOS Press, 1992 pp. 123–137

[8] La Rovere MT et al Baroreflex sensitivity and heart-rate variability in prediction of total cardiac mortality after myocardial infarction The Lancet 1998 351: 478–484

[9] Ferrari GM et al Baroreflex sensitivity predicts long-term cardiovascular mortality after myocardial infarction even in patients with preserved left ventricular function J Am Coll Cardiol 2007; 50: 2285–90

[10] La Rovere MT, Pinna GD, Maestri R, et al Prognostic implications of baroreflex sensitivity in heart failure patients in the beta-blocking era J Am Coll Cardiol 2009; 53: 193–199

[11] La Rovere MT et al Baroreflex sensitivity and heart rate variability in the identification of patients at risk for life-threatening arrhythmias: implications for clinical trials Circulation 2001; 103: 2072–2077

[12] La Rovere MT, Pinna GD, Maestri R, Sleigh P Clinical value of baroreflex sensitivity Netherlands Heart Journal 2013; 21: 61–63

[13] Mathias CJ and Bannister R (Eds) Autonomic Failure: A textbook of clinical disorders of the autonomous nervous system Oxford: Oxford University Press, 2013.

[14] Steptoe A and Vögele C Cardiac baroreflex function during postural change assessed using non-invasive spontaneous sequence analysis in young men Cardiovascular Research 1990; 24: 627–32

[15] Kardos A, Rudas L, Simon J, Gingl Z and Csanady M Effect of postural changes on arterial baroreflex sensitivity assessed by the spontaneous sequence method and Valsalva manoeuvre in healthy subjects Clin Auton Res 1997; 7: 143–148

[16] James MA and Potter JF Orthostatic blood pressure changes and arterial baroreflex sensitivity in elderly subjects Age and Ageing 1999; 28: 522–530

[17] Mattace-Raso F et al Arterial stiffness, cardiovagal baroreflex sensitivity and postural blood pressure changes in older adults: The Rotterdam Study J Hypertens 2007; 25: 1421–1426

[18] Fadel PJ and Raven PB Human investigations into the arterial and cardiopulmonary baroreflexes during exercise Exp Physiol 2011; 97: 39–50

[19] Schwartz CE and Stewart JM The arterial baroreflex resets with orthostasis Front Physiol 2012; 3: 461

[20] Dampney RAL Resetting the baroreflex control of sympathetic vasomotor activity during natural behaviors: Description and conceptual model of central mechanisms Front Neurosci 2017; 11: 461

[21] Holwerda SW et al Arterial baroreflex control of sympathetic nerve activity and heart rate in patients with type 2 diabetes Am J Physiol Heart Circ Physiol 2016; 311: H1170–H1179

[22] Bauer A et al Deceleration capacity of heart rate as a predictor of mortality after myocardial infarction: cohort study. The Lancet 2006; 367: 1674–1681

[23] Bisognano JD, Bakris G, Nadim MK, Sanchez L, Kroon AA, Schafer J, De Leeuw PW and Sica DA Baroreflex activation therapy lowers blood pressure in patients with resistant hypertension: results from the double-blind, randomized, placebo-controlled rheos pivotal trial. J Am Coll Cardiol 2011; 58 : 765–773

[24] Bartsch H, Bartsch C, Mecke D and Lippert TH Seasonality of pineal melatonin production in the rat: possible synchronization by the geomagnetic field Chronobiology international 1994; 11 : 21–26

[25] Vosko AM, Colwell CS and Avidan AY Jet lag syndrome: circadian organization, pathophysiology, and management strategies Nat Sci Sleep 2010; 2 : 187–198

[26] Ramkisoensing A and Meijer JH Synchronization of Biological Clock Neurons by Light and Peripheral Feedback Systems Promotes Circadian Rhythms and Health Front Neurol 2015; 6 : 128

[27] Bartsch RP, Kantelhardt JW, Penzel T and Havlin S Experimental evidence for phase synchronization transitions in the human cardiorespiratory system Phys Rev Lett 2007; 98 : 054102

[28] Bartsch RP, Schumann AY, Kantelhardt JW, Penzel T and Ivanov PCh Phase transitions in physiologic coupling Proc Nat Acad Sci 2012; 109 : 10181–10186

[29] Bartsch RP, Liu KKL, Bashan A and Ivanov PCh Network Physiology: How Organ Systems Dynamically Interact PLoS One 2015; 10 : e0142143

[30] Vinik AI, Maser RE, Mitchell BD and Freeman R Diabetic autonomic neuropathy Diabetes Care 2005; 26 : 1553–1579

[31] Spallone V et al. Toronto Consensus Panel on Diabetic Neuropathy Diabetes Metab Res Rev 2011; 27: 639–653

[32] Brignole M et al. 2004 Guidelines on Management (diagnosis and treatment) of syncope-update 2004. The task force on syncope, European society of Cardiology Europace 2004; 6 : 467–537

[33] Novak P Quantitative Autonomic Testing J Vis Exp 2011; 53: 2502

[34] Markelov OA, Bogachev MI, Mamontov OV, Katinas GS An integrated algorithmic and software solution for biological rhythms analysis: Application to long-term data series 2014 Mechanical Engineering, Automation and Control Systems (MEACS), International Conference on 6986933

[35] Pyko NS, Pyko SA, Markelov OA, Bogachev MI and Mamontov OV Analysis of blood pressure and heart rate synchronization under non-stationary conditions 2014 International Conference on Mechanical Engineering, Automation and Control Systems 6986934

[36] Pyko NS, Pyko SA, Markelov OA and Bogachev MI Systolic blood pressure and pulse intervals synchronization 2015 IEEE NW Russia Young Researchers in Electrical and Electronic Engineering Conference pp. 341–344

[37] Julien C The enigma of Mayer waves: Facts and models Cardiovasc Res 2006; 70 :12–21

[38] Bogachev MI, Markelov OA, Pyko NS and Pyko SA Blood pressure – heart rate synchronization coefficient as a complementary indicator of baroreflex mechanism efficiency 2015 Soft Computing and Measurements (SCM), XVIII International Conference on pp. 173–175

[39] Landau S and Everitt BS A Handbook of Statistical Analyses Using SPSS 1st edition. Boca Raton: Chapman and Hall / CRC, 2003

[40] Draper NR and Smith H Applied Regression Analysis 3rd ed. New York: John Wiley & Sons, 2014

[41] Zolotukhin YV, Markelov OA and Bogachev MI A network-based approach to the analysis of geomagnetic fluctuations 2017 IEEE NW Russia Young Researchers in Electrical and Electronic Engineering Conference pp. 761–764

